# Different plasticity of bud outgrowth at cauline and rosette nodes in *Arabidopsis thaliana*

**DOI:** 10.1101/2021.09.26.461886

**Authors:** Franziska Fichtner, Francois F. Barbier, Stephanie C. Kerr, Caitlin Dudley, Pilar Cubas, Colin Turnbull, Philip B. Brewer, Christine A. Beveridge

## Abstract

Shoot branching is a complex mechanism in which secondary shoots grow from buds that are initiated from meristems established in leaf axils. The model plant *Arabidopsis thaliana* has a rosette leaf growth pattern in the vegetative stage. After flowering initiation, the main stem starts to elongate with the top leaf primordia developing into cauline leaves. Meristems in arabidopsis are initiated in the axils of rosette or cauline leaves, giving rise to rosette or cauline buds, respectively. Plasticity in the process of shoot branching is regulated by resource and nutrient availability as well as by plant hormones. However, few studies have attempted to test whether cauline and rosette branching are subject to the same plasticity. Here, we addressed this question by phenotyping cauline and rosette branching in three arabidopsis ecotypes and several arabidopsis mutants with varied shoot architectures. Our results show that there is no negative correlation between cauline and rosette branch numbers in arabidopsis, demonstrating that there is no trade-off between cauline and rosette bud outgrowth. Through investigation of the altered branching pattern of flowering pathway mutants and arabidopsis ecotypes grown in various photoperiods and light regimes, we further elucidated that the number of cauline branches is closely related to flowering time. The number or rosette branches has an enormous plasticity compared with cauline branches and is influenced by genetic background, flowering time, light intensity and temperature. Our data reveal different plasticity in the regulation of branching at rosette and cauline nodes and promote a framework for future branching analyses.

**One sentence summary:** Different plasticity of branching at cauline and rosette nodes of arabidopsis is revealed through detailed correlative analyses of branching under varied genetic and environmental contexts.

## INTRODUCTION

Shoot architecture is a highly plastic trait of plants, providing them with enormous flexibility to adapt to their environment and be successful when growing in competition with other plants. In seed plants, the main plant body has a primary apical-basal axis that is established during early embryo development. This main axis is defined by the meristem at the shoot apex (SAM) and the root apical meristem at the root tip. Axillary meristems in the shoot incorporate pluripotent stem cells that, as the name suggests, are located in the axils of leaves. These meristems are surrounded by protective leaf primordia that collectively form an axillary bud.

The shoot of arabidopsis, which is monopodial, consists of three different metamers described by Schmitz and Theres (1999). Type 1 metamers consist of a very short internode, a leaf and a bud; these metamers form a rosette. Type 2 metamers consist of an elongated internode, a leaf and a bud, this node being termed a cauline node. Type 3 metamers consist of an intermediate length internode and a floral bud without a subtending leaf that develops at the top of the main shoot and branches. Branches developing from the rosette axillary buds usually produce all three kinds of metamers, while cauline buds produce only type 2 and 3 metamers and lack the rosette-like leaf growth phenotype.

In late flowering mutants or in wild-type arabidopsis plants grown in short days, axillary meristems develop first in the axil of older rosette leaves (Grbić and Bleecker, 2000; Long and Barton, 2000). When these plants start to flower, e.g. by shifting them to long day conditions, the vegetative SAM transforms into an inflorescence meristem which now only initiates floral primordia (Smyth et al., 1990; Hempel and Feldman, 1994). After the transition to flowering, leaf primordia are no longer produced at the SAM. This also coincides with a switch in axillary meristem formation, with axillary meristems now initiating basipetally in the axil of existing leaf primordia (Hempel and Feldman, 1994; Stirnberg et al., 1999; Grbić and Bleecker, 2000; Long and Barton, 2000; Stirnberg et al., 2002). In long-day grown wild-type arabidopsis plants there are no data on the timing of meristem initiation in rosette leaves, however the initiation seems to happen only after the floral transition takes place (Aguilar-Martínez et al., 2007).

As the growth of axillary buds at cauline nodes is induced in a similar basipetal sequence, it was proposed that rosette buds are merely activated as part of this sequence (Stirnberg et al., 1999; Stirnberg et al., 2002; Walker and Bennett, 2018) although this has not been examined directly. One perspective of shoot branching is that plants somehow have an optimal number or amount of branches with their outgrowth being regulated by correlative inhibition even if spread across different nodes, rosette and cauline (Finlayson et al., 2010; Walker and Bennett, 2018). Accordingly, if all branches were considered similar, then arabidopsis plants that produce fewer cauline branches would tend to allow the release and growth of more rosette branches. If this were the case, then cauline and rosette branch growth should be negatively correlated.

There is genetic variation in the balance of cauline and rosette branch numbers in arabidopsis. Compared to wild-type plants, several mutants with increased primary rosette branches (R1) do not show differences in the number of primary cauline branches (C1). These include *branched1* (*brc1*) and *brc2* mutants which lack functional transcription factors that belongs to the TEOSINTE/ CYCLOIDEA/ PROLIFERATING CELL NUCLEAR ANTIGEN FACTOR family and that are repressors of branching (Aguilar-Martinez et al., 2007) and the bushy strigolactone synthesis and signalling *more axillary growth* (*max*) *1* and *max2* mutants (Stirnberg et al., 2002). Particularly in the latter, an acropetal growth pattern was observed in the rosette bud growth after bolting (Stirnberg et al., 2002), contradicting a strictly basipetal activation of branching in arabidopsis (Hempel and Feldman, 1994; Stirnberg et al., 2002).

The shoot branching pattern of the *flowering locus t* (*ft*) mutant is a good example of the potential of plants to differ in the number and position of branches, cauline or rosette. The *ft* mutant flowers much later than wild-type plants and produces more cauline branches, but has almost no rosette branches (Seale et al., 2017; Fichtner et al., 2021b). So, compared to wild-type plants, the *ft* mutant would have fewer branches based on rosette branch number, while it would have an increased number of branches based on the sum of cauline and rosette branches (Seale et al., 2017; Fichtner et al., 2021b). The question that remains to be resolved is whether, for any given genotype, branching at cauline nodes negatively impacts branching at rosette nodes, and vice versa (Walker and Bennett, 2018).

This mechanistic and anatomical consideration of branching is important in the context of hormonal and long-distance signalling. In many plants, the shoot tip inhibits the outgrowth of axillary buds by producing a flow of auxin traveling along the main stem, thereby focusing resources on the main shoot (reviewed in Rameau et al., 2015; Barbier et al., 2017). This phenomenon, called apical dominance, can be alleviated by the removal of the shoot tip, allowing dormant buds to grow out into branches. Auxin is produced in the young leaves at the shoot tip and transported downwards in a basipetal manner (reviewed in Domagalska and Leyser, 2011; Brewer et al., 2013; Barbier et al., 2019). Auxin cannot be transported into axillary buds but regulates branching partly via modulating the levels of two other phytohormones – strigolactones and cytokinins – and partly through auxin export from the bud (reviewed in Domagalska and Leyser, 2011; Wang et al., 2018). Auxin is thought to activate the synthesis of strigolactones (Foo et al., 2005; Brewer et al., 2009) that inhibit bud outgrowth (Gomez-Roldan et al., 2008; Umehara et al., 2008), and inhibit the synthesis of cytokinins (Tanaka et al., 2006; Ferguson and Beveridge, 2009; Shimizu-Sato et al., 2009; Müller and Leyser, 2011) that activate bud outgrowth (Sachs and Thimann, 1967; Chatfield et al., 2000). Strigolactones and cytokinins both partially function via regulating the expression of *BRC1* (Aguilar-Martínez et al., 2007; Martín-Trillo et al., 2011; Braun et al., 2012; Dun et al., 2012, 2013). Shade and PHYTOCHROME B deficiency both contribute to the inhibition of bud outgrowth in arabidopsis and grasses (Finlayson et al., 2010; González-Grandío et al., 2013; González-Grandío and Cubas, 2014). ABA signalling plays an important role in inhibiting bud outgrowth in response to shade, probably acting downstream of BRC1 (González-Grandío et al., 2013; Reddy et al., 2013; González-Grandío and Cubas, 2014; Yao and Finlayson, 2015; González-Grandío et al., 2017). After buds are released from dormancy, they export auxin into the main stem enhancing sustained bud outgrowth (Bennett et al., 2006; Prusinkiewicz et al., 2009; Müller and Leyser, 2011; Brewer et al., 2015; Chabikwa et al., 2019).

Shoot branching is also regulated by resource availability. In addition to affecting auxin levels, the growing shoot tip acts as a strong sink for photoassimilates, suppressing bud outgrowth through sugar deprivation (Mason et al., 2014). Increased sugar availability in buds, for example due to shoot tip removal, not only provides a source of carbon to sustain growth, but also triggers different signals, thereby releasing buds from dormancy (Barbier et al., 2015; Fichtner et al., 2017; Barbier et al., 2021). Plants have developed different signalling pathways involved in sugar sensing, thus allowing plants to adjust their metabolism, growth and development to specific environmental conditions (Li and Sheen, 2016; Fichtner et al., 2021a). Some recent work has highlighted that sugar signalling pathways interact with auxin, strigolactone and cytokinin pathways to promote bud outgrowth (Barbier et al., 2015; Bertheloot et al., 2020; Fichtner et al., 2021b; Salam et al., 2021).

In this study, we sought to test correlative inhibition between the cauline and rosette regions and did so by investigating the extent to which rosette branching in arabidopsis is negatively related to cauline branching under varied genetic and environmental contexts. We achieved wide variation in branching and flowering using three different arabidopsis ecotypes, 25 different arabidopsis mutants impaired in strigolactone, auxin or flowering pathways and a variety of different growth conditions, and we undertook correlation analyses to determine whether cauline and rosette branch numbers were correlated with each other. We further analysed whether cauline and rosette branch growth are correlated with leaf numbers which represent the number of sites of branch development and may also correlate with resource availability. Our study provides a new basis of knowledge for the understanding of shoot architecture regulation in arabidopsis and offers a framework for future branching analyses.

## RESULTS

### Cauline branching and rosette branching show differences in plasticity in response to the environment

To determine whether the number of primary rosette branches (R1) depends on the number of primary cauline branches (C1), we collected phenotypic data obtained in a range of wild type and mutant arabidopsis plants with different shoot architectures (Fig. 1). These included wild type and mutant plants grown in long photoperiods (16-h light) at normal and high planting density. For all experiments, R1 and C1 were scored separately along with rosette leaf number (RL) and cauline leaf number (CL) (Table S1). These data were obtained from five different independent laboratories and therefore also span a range of lab-specific conditions (explained in detail in the Material and Methods section and Table S1). We have therefore presented a number of independent experiments, many of which utilise the same genetic material and similar but not identical growth conditions.

**Figure 1.**
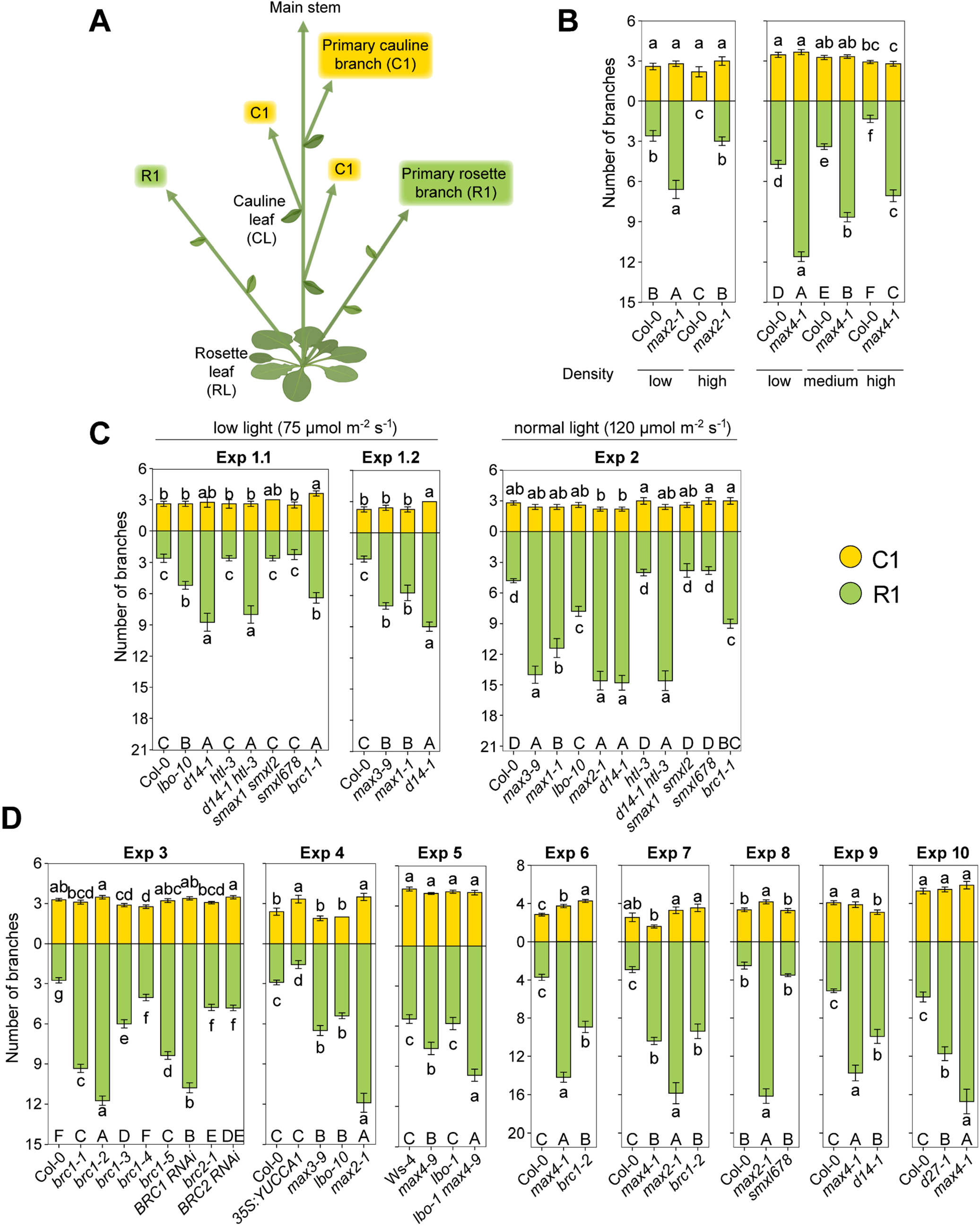
Different variation in cauline and rosette branching occurs in arabidopsis wild type and branching mutant plants. (A) Schematic representation of the arabidopsis branching structure and nomenclature of branching and flowering traits. (B) Arabidopsis wild type (Columbia-0, Col-0) and *max* mutant plants were grown at different planting densities in a 16-h photoperiod. (B) Arabidopsis wild type Col-0 and mutant plants were grown under different light intensities. (D) Wild-type plants (Col-0 or Wassilewskija-4, Ws-4) and branching mutants were grown in 16-h photoperiods. Cauline (yellow, C1) and rosette (green, R1) branch numbers are plotted separately. Letters represent significant differences based on ANOVA with post-hoc LSD testing (*p* < 0.05). Depicted is the mean ± SEM. Small letters represent significant differences for C1 or R1 branches, respectively. Capital letters represent differences in total branch number (C1 + R1).

A wide variation in shoot architecture was observed across the range of branching mutant and wild-type plants and experimental conditions (Fig. 1). The relative differences in R1 number were consistently more varied than the differences in C1 as evidenced from the number of significant differences observed for R1 compared with C1 numbers. Large differences were observed in R1 when wild type or branching mutant plants are grown in low compared to high densities (Fig. 1B). However, there was almost no variation in C1 (Fig. 1B). A comparable trend was observed when comparing wild type and mutant plants that were grown under different light intensities; while there was a lot of variation in R1, C1 and CL numbers did not change (Fig. 1C and Fig. S1C). Similarly, when comparing different mutant plants that show varying degrees of branching grown under normal plant densities and long photoperiods, very little variation in C1 was detected despite a very large variation in R1 (Fig. 1D).

To assess the impact of the number of cauline and rosette branches on the interpretation of the overall branching phenotype in each individual experiment, we separately calculated the significant differences for C1 (small letters, top panels) and R1 (small letters, bottom panels) and for the total primary branches (T1, upper-case letters). In most cases R1 showed the same trend or outcome as T1; however, there were some clear exceptions. In experiment 1 (Exp 1), for example, *brc1-1* plants do not have a significantly different branching phenotype compared to *d14-1* and *d14-1 htl-3* mutants when T1 is calculated, but do have a significantly different phenotype when branching is scored based solely on R1 (Fig. 1D). This is driven by a small, not significantly different increase in C1 in *brc1-1* compared to *d14-1* mutants (Fig. 1D).

### The number of rosette branches does not negatively correlate with the number of cauline branches

The results presented in Fig. 1 indicate that cauline and rosette branch numbers are not strongly correlated with each other. To test this, we performed a correlation analysis between the number of C1 and R1 using the data from Fig. 1 and additional wild type and mutant data that was collected across different laboratories (see Table S1 for all data). We observed a significant but very weak positive correlation for C1 and R1 (r = 0.17, R^2^ = 0.03) (Fig. 2A). When only the wild types were plotted, Ws-4 showed a significant positive correlation between C1 and R1 (r = 0.61, R^2^ = 0.37), while there was no correlation in Col-0 (Fig. 2B). We also analysed the correlation between C1 and R1 in Landsberg *erecta* (Ler) wild-type plants and, similar to Col-0, did not detect any correlation between C1 and R1 (Fig. S1A). The same was also visible when C1 and R1 were plotted for each wild-type and individual experiment sorted in ascending order for the number of C1 branches (Fig. S2); the number of R1 did not show the same trend as the number of C1 further illustrating that there is no negative correlation between C1 and R1 numbers (Fig. S2). We also plotted the correlation of C1 and R1 for mutants in each ecotype background separately (Fig. 2C) and observed a significant positive correlation for mutants in Columbia-0 (Col-0) (r = 0.36, R^2^ = 0.13) but not for mutants in Wassilewskija-4 (Ws-4). These results provide no evidence of a negative correlation between C1 and R1 and therefore do not support the hypothesis of correlative inhibition between cauline and rosette regions in intact plant systems.

**Figure 2.**
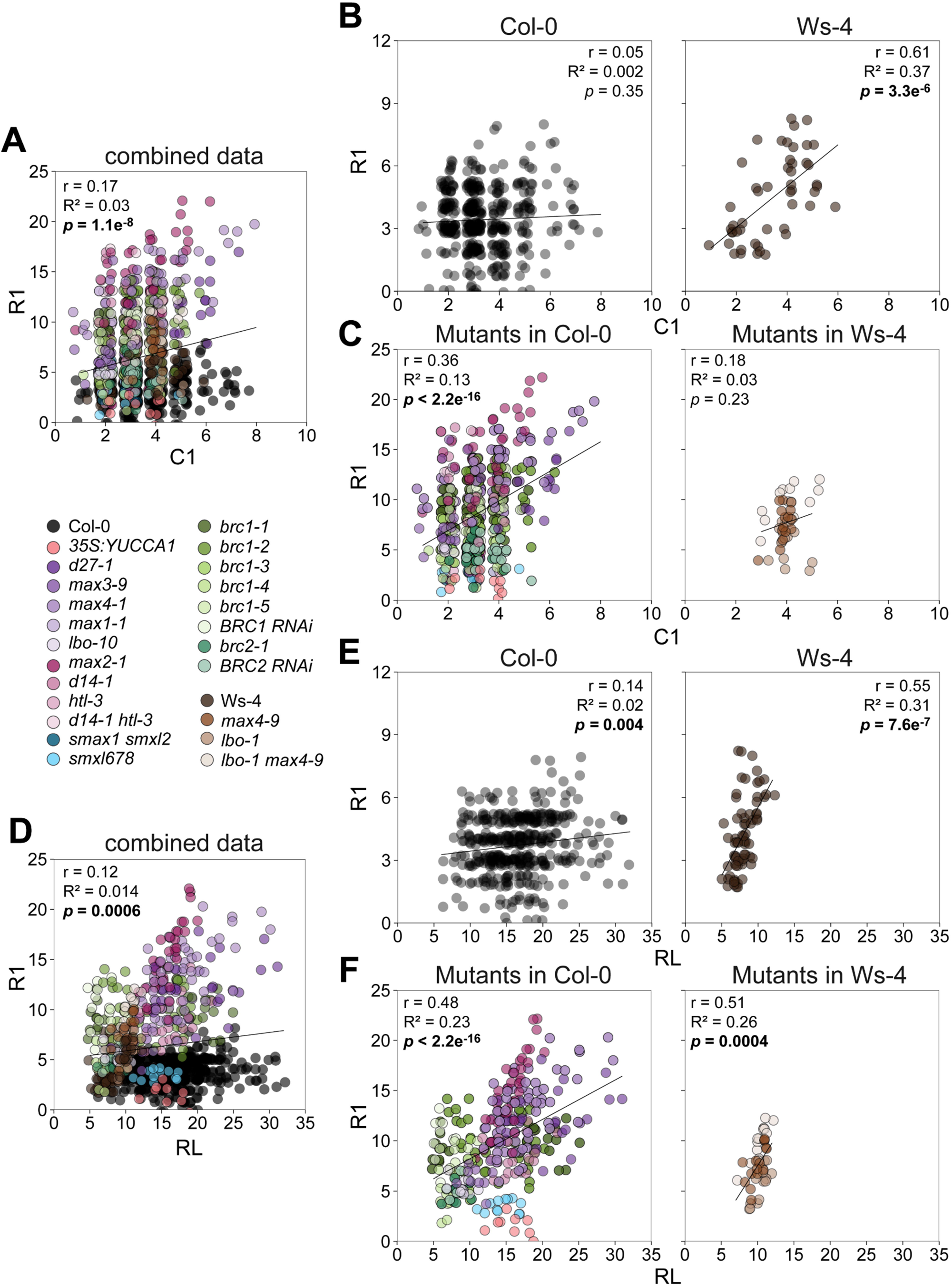
Cauline branch number does not negatively correlate with rosette branch number in arabidopsis wild type and mutant plants. (A-C) The number of cauline branches (C1) or (D-E) the number of rosette leaves (RL) was plotted against the number of rosette branches (R1) and the Pearson correlation coefficient (r), coefficient of determination (R^2^) and probability (*p*) were calculated. (A, D) The correlation for all data presented in Figure 1 and additional data as indicated in Table S1 were used. (B, E) The correlation of C1 and R1 for wild-type plants only. (C, F) The correlation of C1 and R1 in mutant plants was separated by the corresponding ecotype. All plants were grown in a 16-h photoperiod. Genotypes are indicated by different colours. Each data point represents a single plant. Data points were jittered to avoid overplotting and were alpha blended meaning that regions of high point density appear as areas of high colour intensity. Significant correlations are indicated in bold.

### Cauline branch number correlates with cauline leaf numbers while rosette branch number correlates positively with rosette leaf number in arabidopsis mutants with a highly branched phenotype

As the number of nodes of a given metamer type might influence the number of branches produced of that type, we correlated the number of cauline branches and rosette branches with the number of cauline leaves (CL) and rosette leaves (RL) respectively, in a range of arabidopsis lines with different shoot architectures. Using the data from Fig. 1 and additional data (see Table S1 for all data), we observed a strong significant positive correlation (r =0.98, R^2^ = 0.97) between C1 and CL (Fig. S3A). This suggests that under long days, C1 strongly depends on the number of cauline nodes produced (CL). This relationship is upheld when the data were separated by the ecotype (Fig. S3B; Col-0 background: r =0.99, R^2^ = 0.98; Ws-4 background: r =1, R^2^ = 1) or when only wild-type plants were used in the correlation analyses (Fig. S3C; Col-0: r =0.99, R^2^ = 0.97; Ws-4: r =0.91, R^2^ = 0.83; Fig. S1C, *Ler*: r =1, R^2^ = 0.99). In contrast to the strong positive correlation of CL and C1, a very weak but significant positive correlation (r = 0.12, R^2^ = 0.014) was observed between R1 and RL for the combined data set (Fig. 2D). This weak significant positive correlation is maintained in Col-0 plants (Fig. 2E) and branching mutants in the Col-0 ecotype background (Fig. 2F) as well as in Ler wild-type plants (Fig. S1C). Interestingly, wild-type Ws-4 plants (Fig. 2E; r = 0.55, R^2^ = 0.31) and mutants in the Ws-4 background (Fig. 2F; r = 0.51, R^2^ = 0.26) show a stronger significant positive correlation. Consequently, the variation in R1 can be only partly explained by the variation in RL, whereas the variation in C1 is completely related to CL. This shows that there is a different plasticity in cauline and rosette branching and supports a hypothesis whereby cauline and rosette branching may be regulated, at least in part, by different regulatory mechanisms or different emphases within the same regulatory mechanism.

We also correlated C1 and R1 in mutants with different architectures that were grown under at least two different laboratory conditions to ensure enough variability (Fig. 3A, Table S1). In contrast to the *brc1* mutants which did not show a correlation between C1 and R1, a significant strong positive correlation between C1 and R1 was detected in the highly branched strigolactone synthesis and signalling mutants *max4* (r = 0.61, R^2^ = 0.38) and *max2* (r = 0.54, R^2^ = 0.3) (Fig. 3A). We also plotted the correlation between RL and R1 in the same mutants (Fig. 3B). While there was no correlation between RL and R1 in the two *brc1* mutants, a significant strong positive correlation between RL and R1 was detected in *max4* (r = 0.67, R^2^ = 0.45) and *max2* (r = 0.84, R^2^ = 0.71), respectively (Fig. 3B).

**Figure 3.**
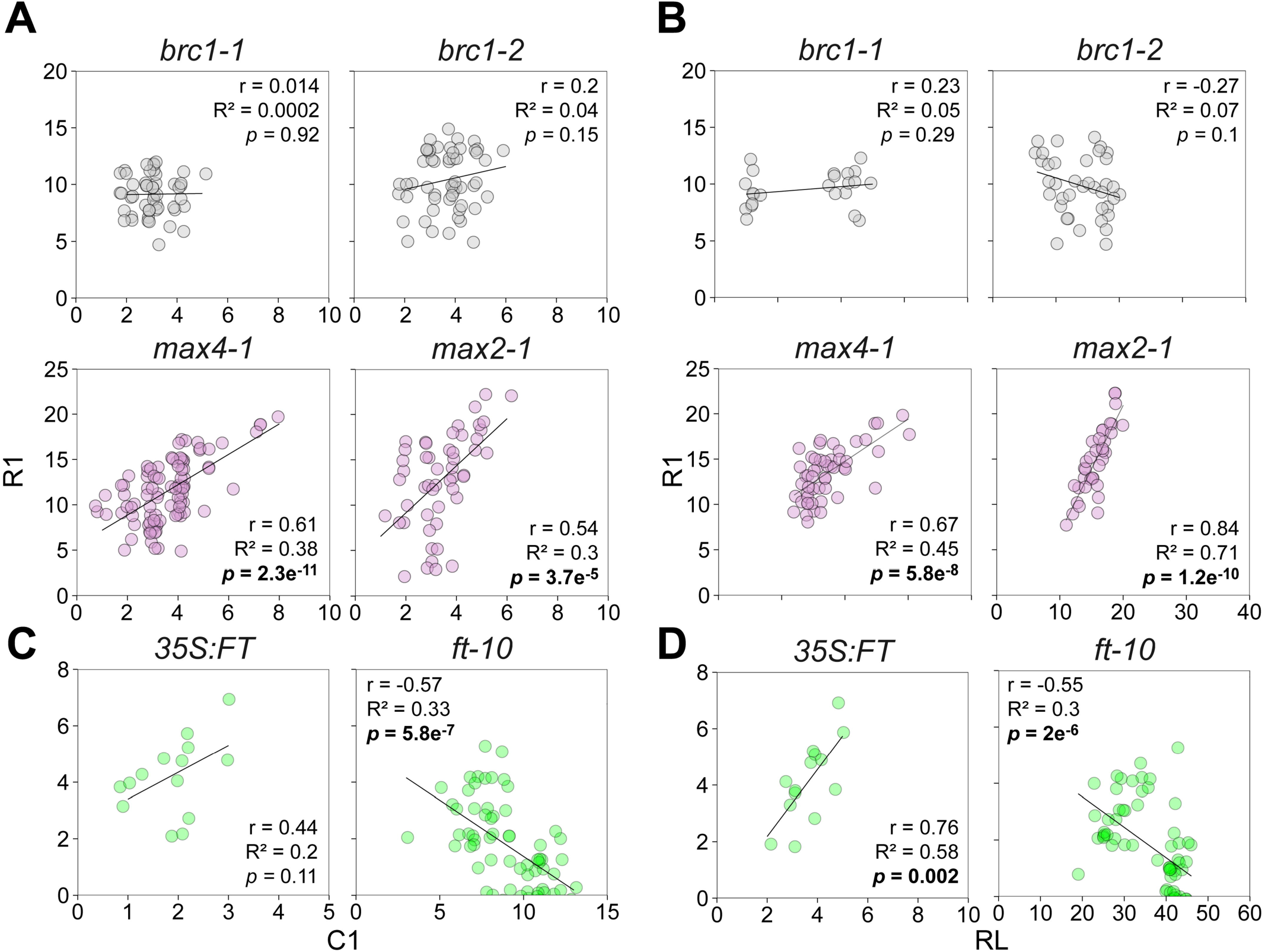
Cauline branch number correlates with rosette branch number in arabidopsis strigolactone and *ft* mutants. (A, C) The number of cauline branches (C1) or (B, D) the number of rosette leaves (RL) was plotted against the number of rosette branches (R1) and the Pearson correlation coefficient (r), coefficient of determination (R^2^) and probability (*p*) were calculated. All mutants were grown in a 16-h photoperiod. Subpanels (A) and (B) were also plotted as part of Fig. 2. All data is presented in the same manner as Fig. 2.

### The flowering pathway is involved in branch outgrowth under long and short-day conditions

As the growth of buds into branches in rosette plants is tied to the bolting stage associated with the flowering process, we explored the relationship between branch growth at cauline and rosette nodes of different flowering lines. The late flowering mutant *ft* is reported to have a strong reduction in rosette branching (Fichtner et al., 2021b) and was compared with a range of other lines affected in flowering time. In contrast to the other mutants analysed, *ft* plants showed a significant negative correlation of C1 and R1 (r = −0.57, R^2^ = 0.33; Fig. 3C). We also plotted the correlation of RL and R1 and observed a significant positive correlation in *35S:FT* (r = 0.76, R^2^ = 0.58) and a significant negative correlation in *ft* mutant plants (r = −0.55, R^2^ = 0.3; Fig. 3D). Summarizing, C1 and R1 did not correlate in wild-type plants or plants with an intermediate branching phenotype (e.g. *brc1* mutants), while C1 and R1 positively corelated in highly branched *max* mutants and negatively correlated in *ft* plants. Additionally, in highly branched plants R1 seems to be highly related to RL, while R1 was less well correlated with RL in plants with an intermediate branching phenotype.

Prompted by these observations, we used flowering mutants and photoperiod to test whether cauline branching and rosette branching are impacted by flowering. Under long days, *ft* and *soc1* mutants have an increase in C1 and a decrease in R1 when compared to Col-0 wild-type plants under the same day-length (Fig. 4A; three independent experiments from two different laboratories). It was previously suggested that this is a consequence of the negative correlation of C1 and R1 branch numbers (Seale et al., 2017). However, our analyses reveal no negative correlation between C1 and R1 in wild-type plants (Fig. 2C). We compared long-day grown *ft* and *soc1* mutants to Col-0 wild-type plants grown in an 8-h photoperiod that show a very similar increase in C1 compared to *ft* and *soc1* mutants (Fig. 4A; two independent experiments from two different laboratories, SD1/ SD2). We did not observe a decrease in R1 in late flowering Col-0 wild-type plants grown in an 8-h photoperiod when compared to those grown in a 16-h photoperiod (Fig. 4A), which was contrary to our observations of late flowering *ft* and *soc1* mutants. This suggests that increased C1 does not necessarily lead to decreased R1; the decreased R1 observed in *ft* and *soc1* mutants is unlikely to be simply due to an increase in C1. We also grew *35S:FT* plants that always have a very early flowering phenotype compared with wild-type plants. These plants have an increase in R1 when compared to Col-0 wild-type plants grown in the same photoperiod (Fig. 4A). We performed an additional experiment with late flowering *ft* and *soc1* mutant plants including another late flowering mutant, *fd*, and grew these plants in increased temperatures to induce earlier flowering to potentially further modulate branching (25/21°C compared to 22/18°C day/night; Fig. 4B). As observed previously (Fig. 4A), all three late flowering mutants have an increase in C1 and a decrease in R1 when compared to Col-0 wild-type plants (Fig. 4B). Interestingly, compared to Col-0 and *ft* plants grown in the same conditions under standard temperatures (22/18°C), both Col-0 wild-type plants and *ft* mutants produce more R1 but the same number of C1 when grown under increased temperatures (Fig. 4B). To further explore the effect of *ft*-mediated flowering and branching, we grew Col-0 wild-type plants and *ft* mutants in short-day conditions where these genotypes produce the same amount of rosette leaves (Col-0 65.8±1.5, *ft* 67.2±1.6, p>0.05; 8-h photoperiod; Fig. 4C). However as observed in long-day conditions, *ft* mutants still developed more C1 and less R1 branches compared to wild-type plants. Consequently, the *ft-*mediated flowering pathway does influence branching in arabidopsis under long and short-day conditions, with high levels of *FT* promoting rosette branching, and low levels of *FT* or downstream signalling components (like *SOC1*) inhibiting it (Fig. 4A-C).

**Figure 4.**
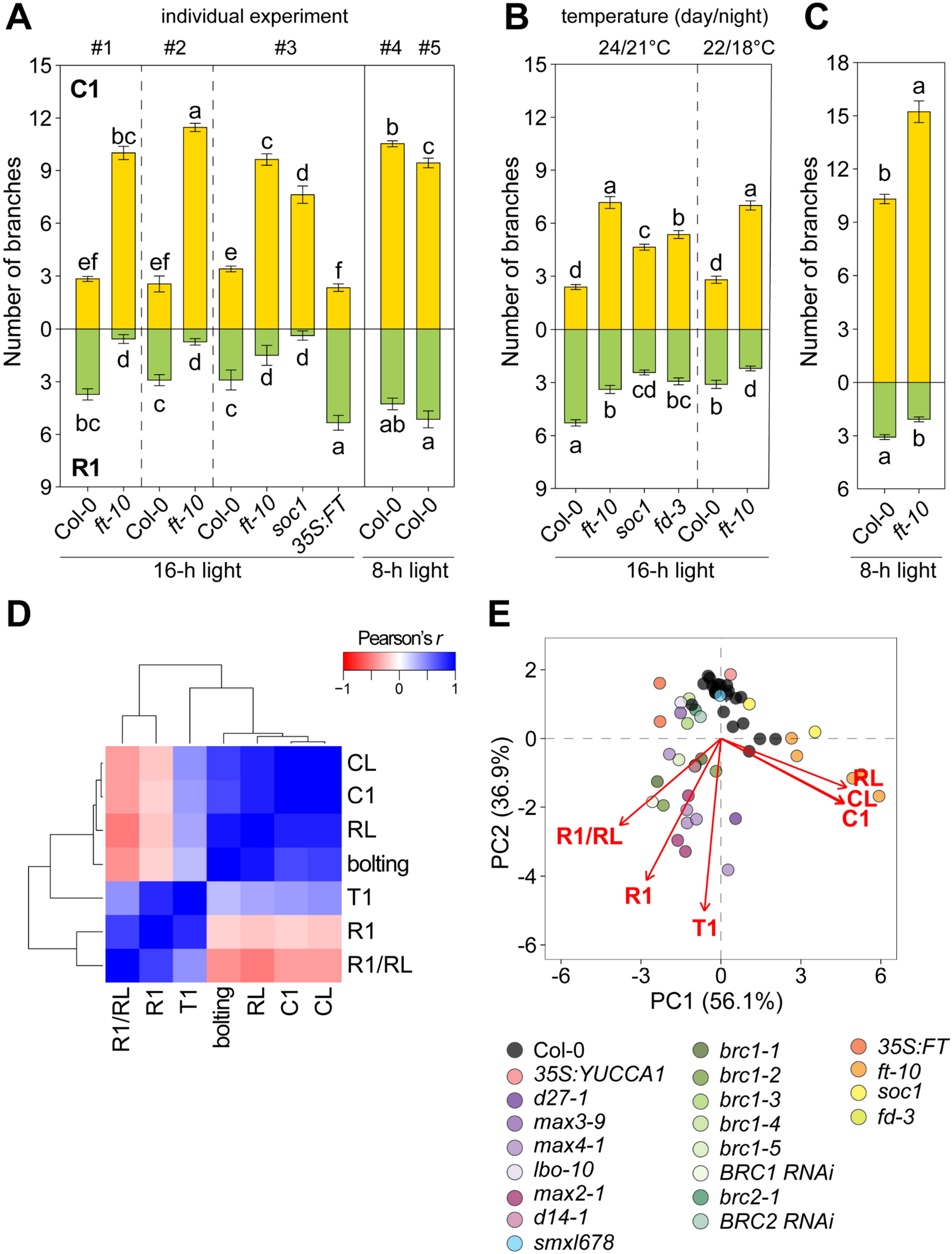
The flowering pathway affects rosette and cauline branch numbers. (A) Arabidopsis Col-0 wild type and flowering mutant plants were grown in 16-h or 8-h photoperiods and the number of primary cauline (C1, yellow) and rosette (R1, green) branches were determined. (B) Col-0 and flowering mutant plants were grown in a 16-h photoperiod under different temperatures and C1 as well as R1 were determined. (C) Col-0 and *flowering locus t* (*ft*) mutant plants were grown in an 8-h photoperiod and C1 as well as R1 were determined. Letters represent significant differences based on ANOVA with post-hoc LSD testing (*p* < 0.05). Depicted is the mean ± SEM. (D) Cluster analysis of branching and flowering traits in Col-0 wild type and mutant plants in the Col-0 background based on the Pearson correlation coefficients (r). Dendrograms represent clusters using a canberra distance matrix with average-based clustering. (E) Principal component (PC) analysis of branching and flowering traits in arabidopsis Col-0 wild type and mutant plants grown in a 16-h photoperiod. Mean values for each trait were used for the analysis. Data points represent a single experiment and were alpha blended meaning that regions of high point density appear as areas of high colour intensity. The percentages of total variance represented by PC 1 and PC 2 are shown in parentheses. The loadings of individual traits are indicated (red). Different colours represent the different genotypes. T1, total number of primary branches; CL, cauline leaf number; RL, rosette leaf number; bolting, days to bolting.

### Cauline branch number clusters with flowering traits in arabidopsis wild type and mutant plants grown under long days

To further investigate the relationship between cauline branching, rosette branching and flowering, as well as to highlight potential mechanisms regulating these processes, a cluster analysis was performed (Fig. 4D). We used the Pearson correlation coefficient of all data available for mutants in the Col-0 background and Col-0 wild-type plants with the following variables: days to bolting (bolting), cauline leaf number (CL), rosette leaf number (RL), cauline branches (C1), rosette branches (R1), total branches (T1, the sum of C1 and R1), and R1 divided by RL (R1/RL) (Fig. 4D, see Table S1 for data set). Hierarchical clustering of the Pearson’s r of these variables led to the formation of two main clusters, with the first cluster comprising CL, C1, RL, days to bolting and T1, and the second cluster comprising R1 and R1/RL (Fig. 4D). Additionally, principal component analysis (PCA) was performed based on the averages of all variables for mutants in the Col-0 background and Col-0 wild-type plants grown in long-day conditions (Fig. 4E; see Table S1 for all data). Here, the strigolactone mutants, the *brc1* mutants and the *35S:FT* line seemed to separate from the Col-0 wild types. Similarly, the late flowering mutants also diverged away from Col-0 wild-type plants (Fig. 4E). Again, these divergences support the notion of independent genetic regulation of the values of flowering related traits (RL, CL, C1) compared with the values of the rosette branching (R1) and related traits T1 and R1/RL. As in the previous cluster analysis based on the Pearson’s r (Fig. 4D), C1, CL and RL were tightly aligned in the PCA as represented by their loading (i.e. the weight they have in the analysis; red arrows on the horizontal axis, Fig. 4D) driving the data along PC1 (56.1% of variation). This is suggesting these traits are highly correlated. Interestingly, the loading for T1 was in the middle of the R1 traits and the highly connected C1/ CL/ RL group (Fig. 4D). T1 and R1 largely drove data separation along PC2, which accounted for most (36.9%) of the remaining variation (Fig. 4D). In conclusion, both approaches (Fig. 4) support the results of the visual inspection (Fig. 1) as well as correlation analyses (Fig. 2, 3) that C1 does not negatively correlate with R1. In addition, the clustering of C1 with flowering dependent traits like leaf number and days to bolting, indicates that C1 might be connected to flowering time.

Interestingly, dividing R1 by RL further separated the data in the PCA. Consequently, R1/RL may be useful to account for variation in the branching phenotype among individuals and between genotypes with altered flowering time and/or leaf number. This may be particularly useful where variation in branching due to environmental effects on flowering are to be minimised.

There is a significant weak positive correlation of RL and R1 in long-day grown wild-type plants, a strong positive correlation of RL and R1 in strigolactone mutants and a range of individual *35S:FT* plants, and a strong negative correlation in *ft* mutants (Fig. 3). This indicates that the number of R1 is somewhat related to RL and thus rosette node numbers in arabidopsis. Therefore, we replotted the data for Fig. 4A and 4B based on R1/RL (Fig. 5A, 5B). This highlighted that, relative to their RL, *ft* and *soc1* plants branch much less when compared to Col-0 plants (grown in either long or short days) independent of the growth temperature and that *35S:FT* plants have a strong increase in branching at rosette nodes (Fig. 5A, 5B). Interestingly, *35S:FT* plants seem to produce more than one R1 per rosette leaf, indicating that, similar to strigolactone mutant plants, they are likely limited in branching by the number of leaves/nodes developed. This would also explain why *35S:FT* mutants seemed to cluster with strigolactone mutants in the PCA (Fig. 4D). In order to compare branching in *35S:FT* and *max* mutants, we subsequently plotted R1/RL for all available experiments with *max4* and *max2* plants and compared them to available experiments with *35S:FT* plants. While only a minor increase in R1 was detected in *35S:FT* plants compared to wild-type controls (Fig. 4A), from the perspective of R1/RL, *35S:FT* plants actually branch to a similar degree as the *max4* and *max2* mutants (Fig. 5C).

**Figure 5.**
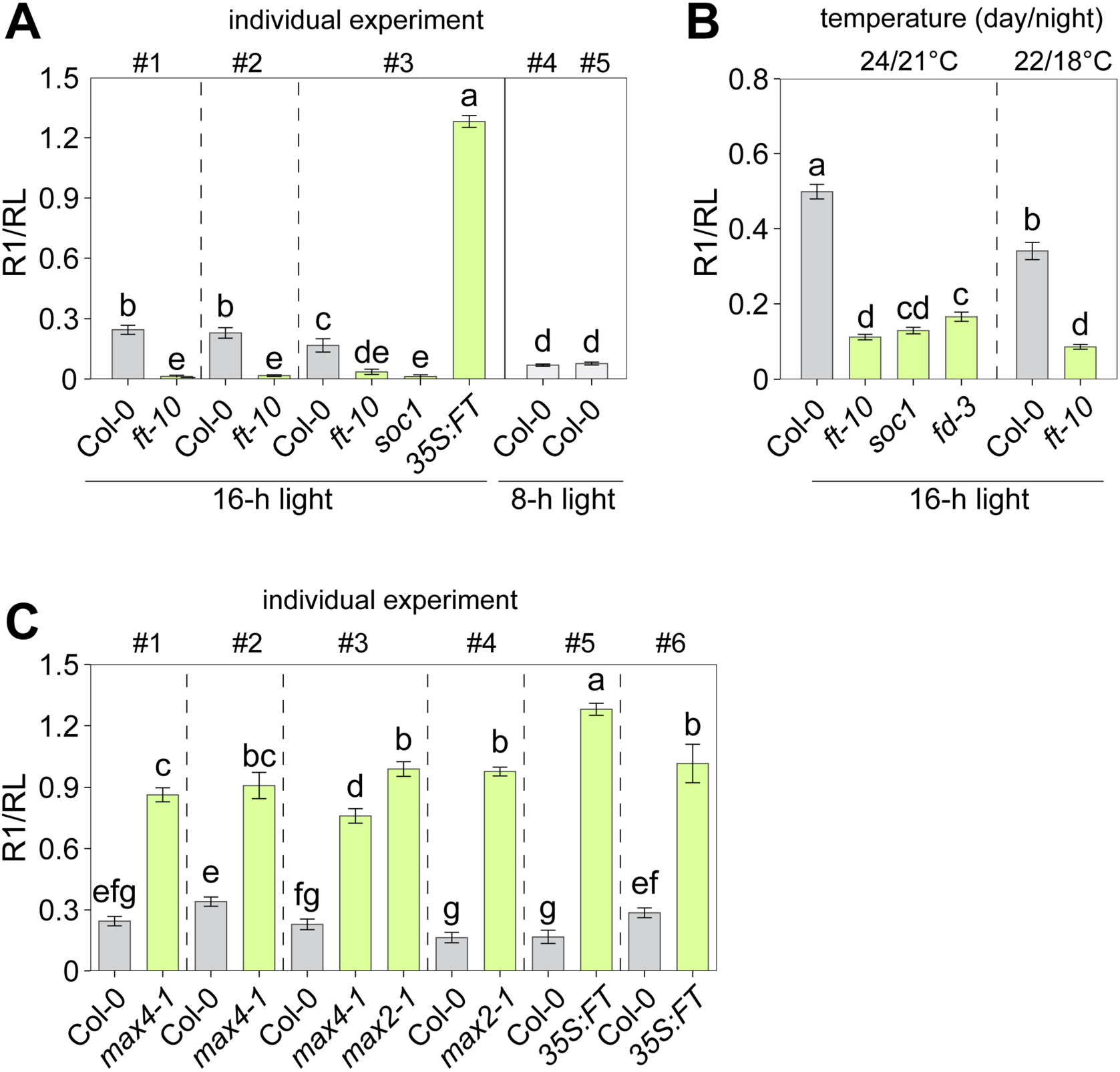
Exemplar where rosette leaf number can be used to normalize rosette branch number in highly branched plants. (A, B) Data from Figure 4A and 4B was replotted based on the number of rosette branches per rosette leaves (R1/RL). (C) The number of R1/RL in highly branched (green) and Col-0 wild-type plants (grey) are shown. Letters represent significant differences based on ANOVA with post-hoc LSD testing (*p* < 0.05). Depicted is the mean ± SEM.

### Cauline branch number consistently correlated with flowering time in different arabidopsis ecotypes and photoperiods

To further investigate the relationship between C1, R1 and flowering time measures, we sought to increase the variability in C1, R1, CL and RL numbers by investigating three arabidopsis ecotypes grown in a variety of photoperiods and light intensities (Fig. 6A). In Col-0, Ler and Ws-4 wild-type plants, C1, CL, and RL consistently increased in shorter photoperiods (a single experiment is given as an example in Fig.S4A-C; all data can be found in Table S1). In contrast, R1 was less related to photoperiod and more related to light regime (Fig. S4A-C). This further supports the hypothesis that cauline and rosette buds have a different response to environmental and endogenous signals and are therefore not regulated equivalently.

**Figure 6.**
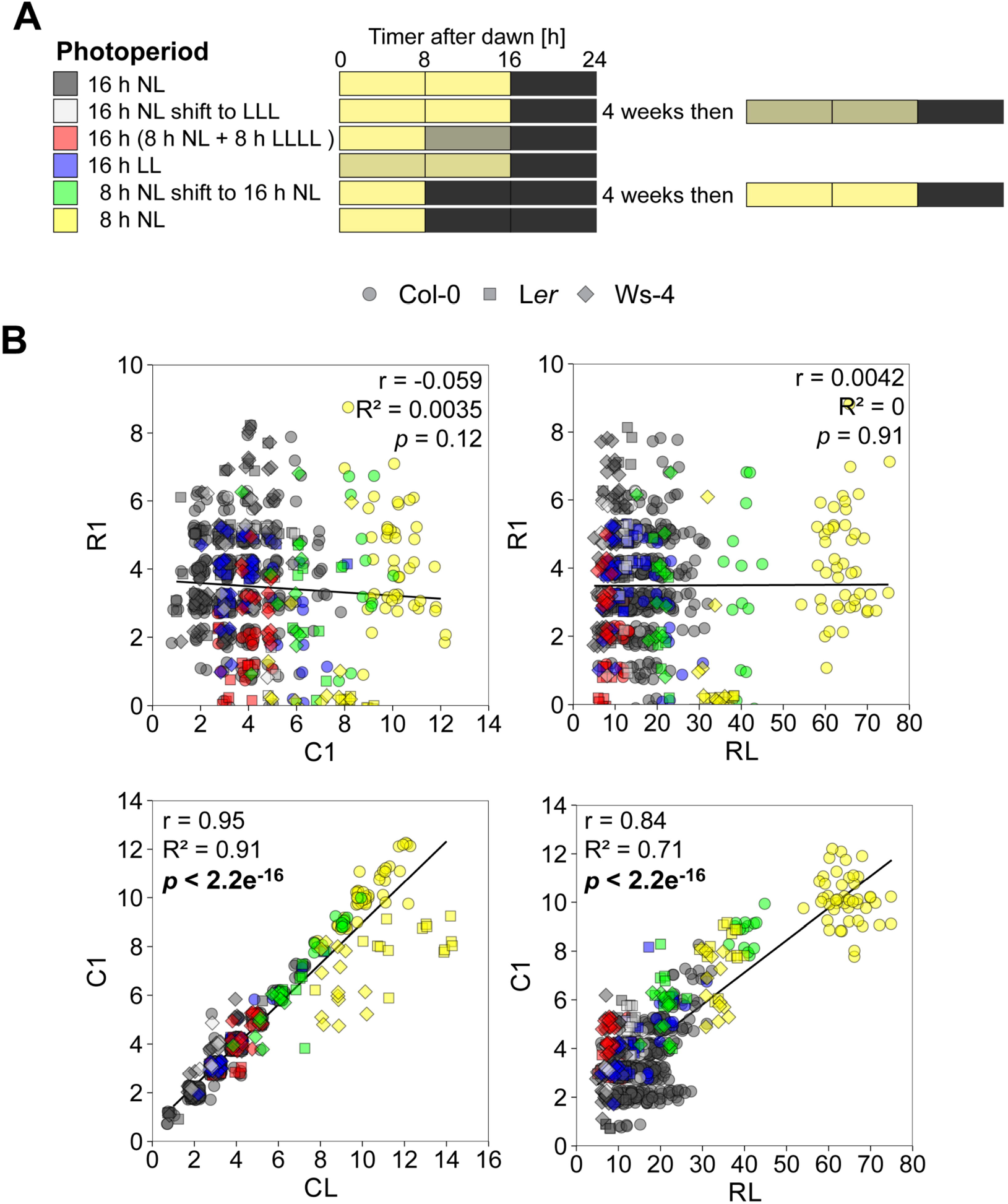
Cauline branch number is not correlated with rosette branch number in arabidopsis wild-type plants grown under different photoperiods. (A) Arabidopsis ecotypes (Columbia-0, Col-0 (circles); Landsberg *erecta*, Ler (squares); Wassilewskija-4, Ws-4 (diamonds)) were grown in a variety of photoperiods. Different shades of yellow to black represent different light intensities; bright yellow for 150 μmol m^-2^ s^-1^ (NL), to decreasing light intensities of 75 μmol m^-2^ s^-1^ (LL), 40 μmol m^-2^ s^-1^ (LLL) and 5 μmol m^-2^ s^-1^ (LLLL) with all plants experiencing at least 8 h of complete darkness (shown in black). (B) Correlation analyses of the number of primary cauline and rosette branches (C1 and R1, respectively) and leaf numbers (CL and RL, respectively). The Pearson correlation coefficient (r), coefficient of determination (R^2^) and probability (*p*) were calculated. Each data point represents a single plant. Data points were jittered to avoid overplotting and are alpha blended. Significant correlations are indicated in bold.

Correlation analysis using the combined available data from these different ecotypes (including data from Fig. 1-4) confirmed our previous results: no correlation was obtained between the number of C1 and R1, but a significant positive correlation was observed for the number of C1 and CL (Fig. 6B). No correlation between RL and R1 was detected (Fig. 6B). However, a strong positive correlation between C1 and RL was detected in all three ecotypes (Fig. 6B), as these traits largely depend on flowering time. The same relationships were detected when the data were correlated based on the mean values for each individual experiment (Fig. S4D).

When examining the correlation of C1 and CL, *Ler* and Ws-4 ecotypes grown under short day conditions diverted from the linear relationship of C1 and CL (Fig. 6B, S5D) implying that not all cauline leaf axillary buds elongated in these ecotypes under this photoperiod. Interestingly, under short day or short-day to long-day shift conditions, lower node cauline branches could elongate before upper node cauline branches (Fig. S5A-B) and, in some instances, rosette branches grew out before upper node cauline branches were activated (Fig. S5C-D).

To investigate if these effects are simply due to photoperiod effects, we also performed correlation analyses for all three ecotypes in long-day grown plants only (Fig. S6A). The results are very similar to the analyses done on the combined photoperiod data set: high positive correlations between CL and C1 as well as RL and C1; no correlation between RL and R1 (Fig. S6A). In long day conditions, however, a significant positive correlation between C1 and R1 was obtained, although with a very low R^2^ of 0.01 indicative of a very weak and potentially not biologically relevant correlation (Fig. 5A).

To investigate the relationships between C1, R1, days to bolting, CL and RL in highly branched plants, a cluster analysis based on the Pearson’s r was performed on the data from Fig. 5 (Fig. 7A). The results were remarkably reminiscent of those obtained using the set of mutants grown in long-day conditions (Fig. 4D). Hierarchical clustering led to the formation of two main clusters: R1 and R1/RL formed one cluster, and days to bolting, RL, C1, CL, and T1 formed the other cluster (Fig. 7A). The same clustering was obtained when only the experiments of long-day grown ecotypes were analysed (Fig. S6B). We also performed PC analyses using the mean of each experiment, which again gave very similar results (Fig 6B). C1, RL and CL drove the data in the same direction along PC 1 (63.6% of the total variation). R1 on the other hand drove the data along PC 2 (29.3% of the total variation). The loading of T1 was between R1 and the highly linked group of C1, CL, RL. In contrast, R1/RL drove the data in the other direction in an orthogonal way, further separating it (Fig. 6B). These analyses in different arabidopsis ecotypes (Fig. 7B) and different mutants (Fig. 4E) suggest that dependent on the biological question, using T1 (R1 + C1) as a parameter for branching may be inappropriate, especially in plants that flower at different times (Fig 8). In contrast, R1/RL facilitates some correction of the data for differences in flowering time that are tightly linked to differences in leaf number (Fig. 8).

**Figure 7.**
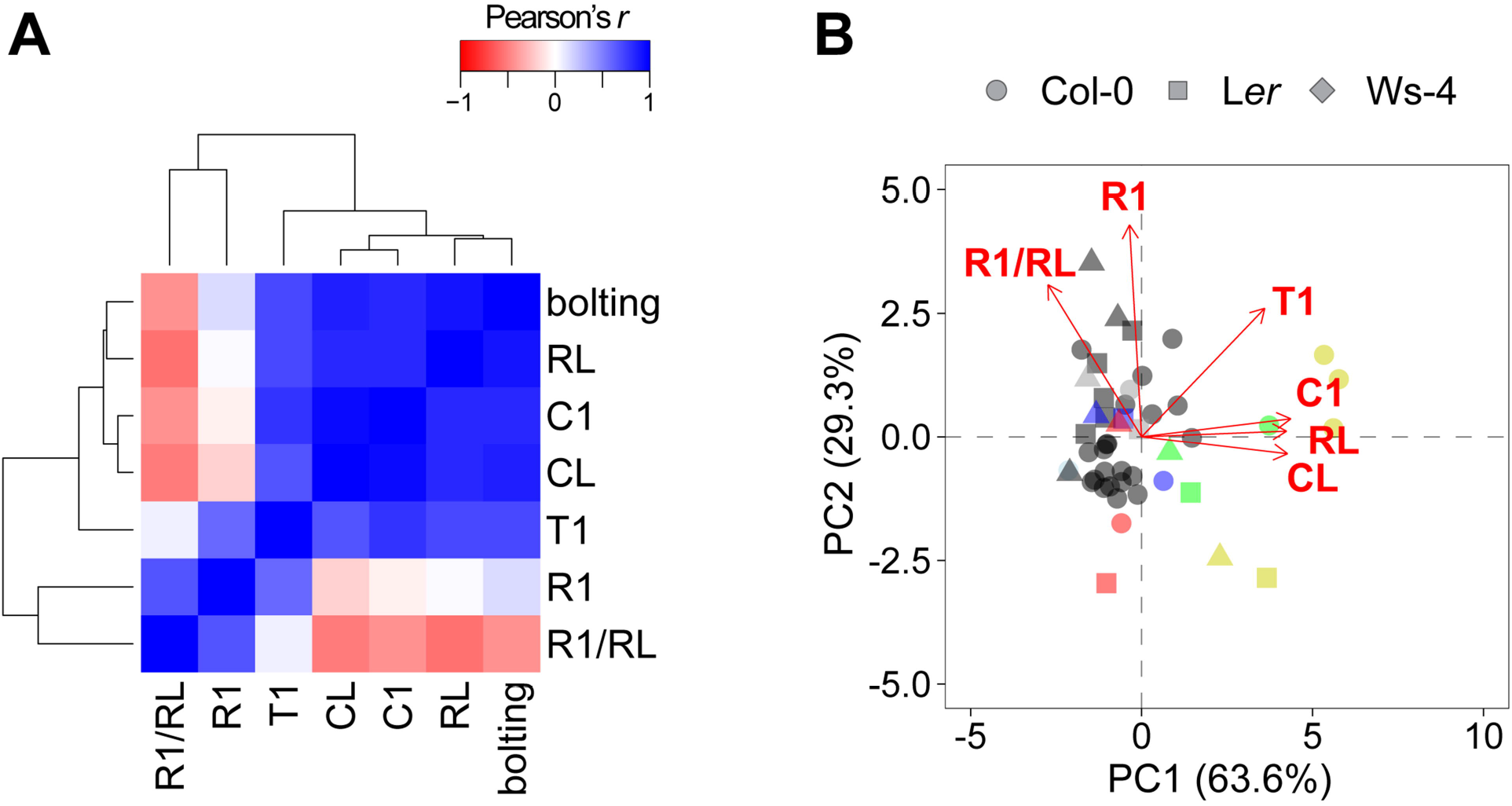
Cauline branch number clusters with flowering traits. (A) Cluster analysis of branching and flowering traits based on the Pearson correlation coefficients (r) in different arabidopsis ecotypes. Dendrograms represent clusters using a canberra distance matrix with average-based clustering. (B) Principal component (PC) analysis of branching and flowering traits in different arabidopsis ecotypes (Columbia-0, Col-0 (circles); Landsberg *erecta*, Ler (squares); Wassilewskija-4, Ws-4 (triangles)). Mean values for each trait were used for the analysis. Data points represent a single experiment and were alpha blended (regions of high point density show up as areas of high colour intensity). The percentages of total variance represented by PC 1 and PC 2 are shown in parentheses. The loadings of individual traits are indicated (red). Different colours represent the different photoperiods as indicated in Fig. 6A. R1, rosette branch number; T1, total number of primary branches; CL, cauline leaf number; C1, cauline branch number; RL, rosette leaf number; bolting, days to bolting.

**Figure 8.**
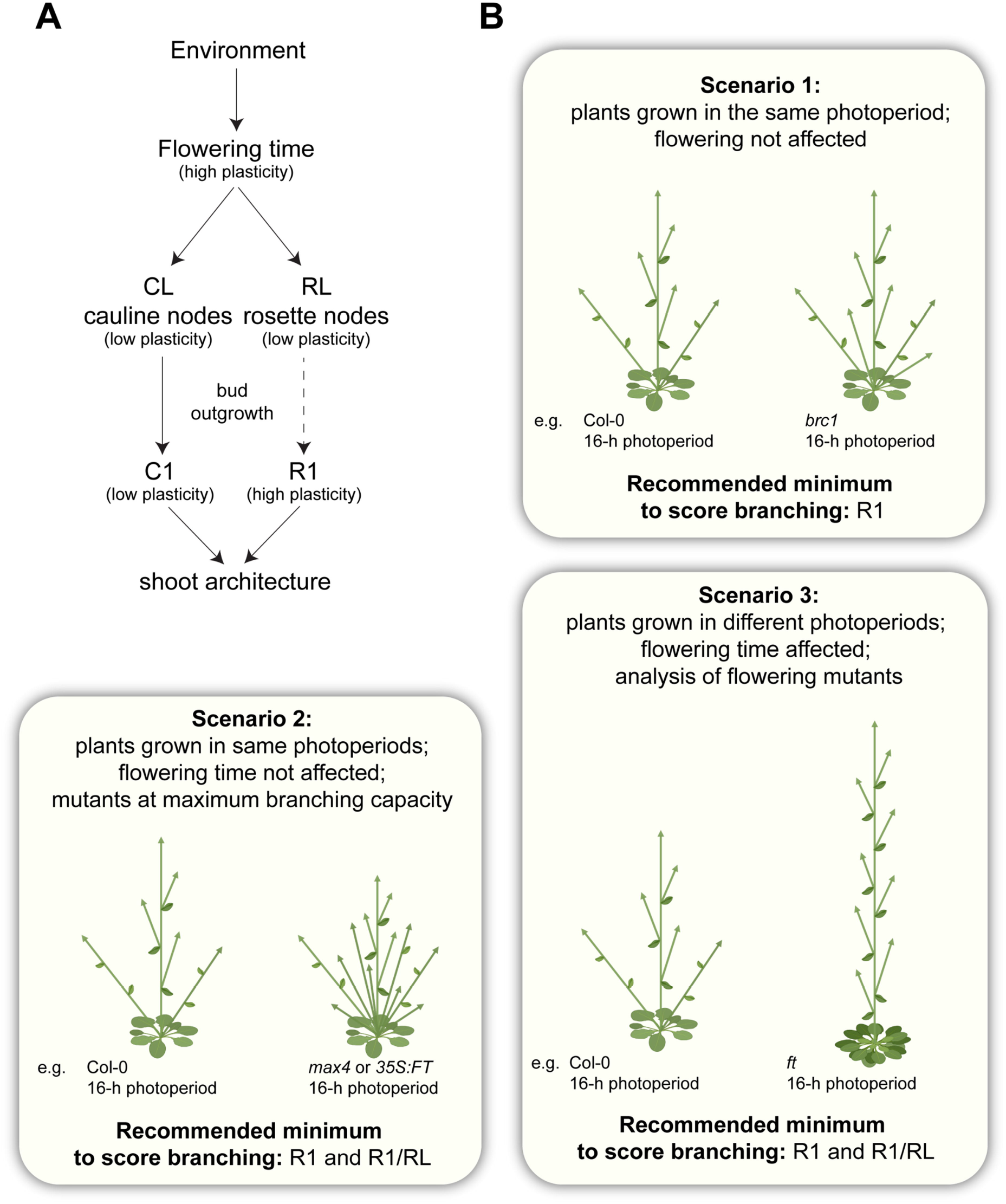
Representation of the architectural plasticity of arabidopsis shoot branching and scenarios showing implications for analysis of rosette branching in arabidopsis. (A) Environmental factors influence flowering time which in turn directly influences cauline leaf (CL) and rosette leaf (RL) numbers and therefore influences the respective node numbers. There is low plasticity in C1 branch outgrowth as the number of CL directly influences the amount of cauline branches (C1) (correlation ≈ 1). There is high plasticity in R1 branch outgrowth as the number of RL only partially influences the number of rosette branches (R1) in wild-type plants. Both C1 and R1 numbers determine the final shoot architecture. The dashed line represents partial dependency. (B) Scenario 1 compares arabidopsis plants that are grown in the same photoperiod and that have the same flowering time and the same RL but are not extremely bushy. In this scenario, R1 can be counted as a representation of the rosette branching phenotype. Scenario 2 is the same as Scenario 1, but includes plants that are close to their maximum branching capacity (i.e. close to 1 R1 branch per rosette leaf or node). In this scenario, R1 as well as R1/RL should be analysed due to the impact of any variation in leaf number on R1. For Scenario 3, where plants differ in flowering time and therefore RL, again both R1 as well as R1/RL should be analysed, and attention given to the impact of flowering on RL and cauline branch number. Images are intended as schematic representations.

## DISCUSSION

### Branching at cauline and rosette nodes are independent variables subject to different developmental plasticity

In this study, we showed that R1 and C1 are not negatively correlated in arabidopsis wild type and mutant plants providing little evidence for correlative inhibition between the cauline and rosette regions in intact plants. As there is no negative trade-off between these variables, branching at rosette and cauline nodes highlights potential differences in, for example, gene regulatory, hormonal and/or environmental variables during ontogenetic development in arabidopsis. As such, C1 and R1 should be treated separately. We showed that C1 is highly correlated with the number of cauline nodes (CL) produced across our wide experimental range (Fig. 6, S1, S3, S4, S6). Our study highlights that, contrary to cauline branching, the variation in the number of rosette nodes only partly affects the number of rosette branches in wild-type plants (Fig. 8A). So, while there is only limited plasticity of branch outgrowth at cauline nodes, there is an enormous plasticity at rosette nodes, suggesting that there must be certain differences in their outgrowth regulation. However, in highly branched mutants (*max4, max2*), the number of rosette branches is highly correlated with the number of rosette leaves. A significant correlation was also observed in plants overexpressing *FT* and which are very early flowering. Using clustering analyses, we demonstrated that C1 clusters with traits related to flowering time (RL and bolting date). This explains the positive correlation of R1 and C1 as well as of RL and R1 in mutants that branch at their maximum capacity (R1/RL is close to 1 in *max4, max2, 35S:FT*, Fig. 5) where the limiting factor of branching is the number of nodes produced (reflected by the number of rosette leaves). When highly branched plants flower late, they develop more RL and CL, leading to the formation of more R1 and C1. As these plants develop branches at almost every node, this would in turn cause the positive correlation between C1 and R1 in these genotypes. Thus, when comparing the branching phenotype of plants affected in flowering and/or in the number of nodes produced, dividing R1 by RL will partially account for differences in RL and thus better highlight significant effects (Fig. 8B). Moreover, as C1 and R1 are shown here to be not negatively correlated and probably not part of the same/dependent activation sequence, the total number of primary branches should not be used to assess branching phenotypes in arabidopsis as this obfuscates the branching phenotypes. This is also highlighted by how T1 influenced the PCs (Fig. 4E, 7B). Instead R1 and C1 should always be stated separately (Fig. 8B; Aguilar-Martínez et al., 2007; González-Grandío et al., 2013; Chevalier et al., 2014; Brewer et al., 2016; González-Grandío et al., 2017; Barbier et al., 2021; Fichtner et al., 2021b). This is of special importance when plants flower at different time points or are grown under different photoperiods.

We also detected a weak but significant positive correlation of RL and R1 in Col-0, Ws-4 and Ler wild-type plants (Fig. 2F, S1B). This suggests that, in wild-type arabidopsis, part of the differences in R1 depends on RL number. The correlation between leaf number and branching in long-day conditions might be a consequence of increased sugar supply as more leaves would usually produce more total photoassimilates. Evidence that carbon/sugar availability influences R1 development in arabidopsis has been obtained with plants grown in low light conditions or exposed to a night extension. Barbier *et al*. (2021) quantified the very early rosette bud growth that occurs after the floral transition but before bolting in long days, and observed less growth in plants with less photosynthetically active light. Recent advances in shoot branching research have illustrated that the release of bud dormancy and outgrowth into a new branch are dependent on sugar availability and involve sugar signalling pathways, notably mediated by trehalose 6-phosphate (Tre6P) and HEXOKINASE1, which interact with the hormones controlling branching (Mason et al., 2014; Barbier et al., 2015; Fichtner et al., 2017; Tarancón et al., 2017; Bertheloot et al., 2020; Barbier et al., 2021; Fichtner et al., 2021b; Salam et al., 2021).

### The FT-mediated flowering pathway is involved in rosette branch regulation in arabidopsis

In arabidopsis, FT is synthesized in phloem companion cells in leaves under inductive long-day conditions and moves in the phloem sieve elements to the SAM, where it interacts with the FLOWERING LOCUS D protein to promote floral transition (Turck et al., 2008). There is a growing body of evidence suggesting that FT is an important regulator of branching, based on studies in rice (*Oryza sativa;* Tsuji et al. (2015)), tomato (*Solanum lycopersicum;* Weng et al. (2016)) tobacco, (*Nicotiana tabacum;* Li et al. (2015)), and pea (Beveridge and Murfet (1996); Hecht et al. (2011)). Flowering in arabidopsis is dependent on Tre6P synthesis (Schluepmann et al., 2003; Wahl et al., 2013; Fichtner et al., 2020), a sucrose specific signalling metabolite in plants (Fichtner and Lunn, 2021). *FT* transcription is also a target of Tre6P signalling (Fichtner et al., 2021b). Plants with higher Tre6P in the vasculature have an early flowering and an increased branching phenotype, and this coincides with upregulation of *FT* (Fichtner et al., 2021b). Stimulation of branching by increased Tre6P in the vasculature was abolished in an *ft* mutant background (Fichtner et al., 2021b), further implicating FT in the regulation of branching in arabidopsis.

Here, we showed that wild-type Col-0 plants that have an increase in C1 similar to the level observed in *ft* plants, still initiate R1 and do not have a decreased R1, unlike *ft* plants. However, in contrast to wild-type plants, C1 and R1 were negatively correlated in *ft* mutants as were RL and R1 (Fig. 3C, D). This affirms two of the observations we made previously, that C1 branch number is tightly related to flowering and that the flowering pathway is also involved in R1 branch number regulation. Triggering earlier flowering via, for example, an increase in temperature, also increased the number of R1 in wild-type and *ft* mutant plants (Fig. 4B). However, the number of R1 in *ft* mutants was always lower than the respective number in wild-type plants grown under the same temperatures, demonstrating that the FT-mediated flowering pathway is involved in regulating rosette branching in arabidopsis. We further showed that the FT-mediated flowering pathway also seems to be important for rosette branch outgrowth regulation in short-day conditions as *ft* mutants also produce less R1 branches compared to Col-0 wild-type plants in short days (Fig. 4C). It was demonstrated previously that there is detectable *FT* expression in short days especially under elevated ambient temperatures, although much lower when compared to long-day conditions (Yamaguchi et al., 2005; Balasubramanian et al., 2006; Lee et al., 2007; Kim et al., 2012). This builds on our speculation that FT has a role for bud outgrowth in short and long-day conditions.

It has been demonstrated that FT can move not only to the SAM but also to axillary meristems, and promote their elongation and development (Niwa et al., 2013; Tsuji et al., 2015; Dixon et al., 2018). FT in arabidopsis, wheat, and hybrid aspen has been shown to interact directly with BRC1, and this interaction leads to a reciprocal repressive effect between the two proteins (Niwa et al., 2013; Dixon et al., 2018; Maurya et al., 2020). *35S:FT* plants developed more than one R1 per RL (Fig. 4C). This is very similar to the phenotype of *brc1* mutants that overexpress a Tre6P synthase in the vasculature resulting in an increase in Tre6P (Fichtner et al., 2021b). We speculated that this phenotype in *brc1* plants with high levels of Tre6P might be a consequence of higher levels of FT and the loss of BRC1 having a strong additive effect on branching (Fichtner et al., 2021b). This would also be a plausible explanation for the branching phenotype of the *35S:FT* plants that have potentially a very strong and relatively constitutive overexpression of *FT*, so potentially a complete inhibition of BRC1 activity, resulting in bud release.

### Shoot branching regulation in arabidopsis rosette and cauline nodes is influenced differently by photoperiod and light intensity

By growing different arabidopsis ecotypes in a wide variety of different light regimes and photoperiods, we demonstrated that C1 is influenced by flowering time, while R1 seemed to be more related to the light regime and intensity (Fig. 6, 7, 8A). While we detected significant positive correlations between RL and R1 in all three ecotypes when the relationship was analysed in long-day conditions and separated by ecotype, there was no correlation when data from different photoperiods was combined (Fig. 6B) or when data from long days and all three ecotypes was merged (Fig. S6A). This provides evidence of genetic regulation of the relation of RL to R1. Interestingly, the correlation between RL and R1 seems to be stronger in ecotypes that develop less RLs as the correlation in Ws-4 is much stronger compared to Col-0 (Fig. 2F). This is in line with sugars having an important role in rosette branching. It would be interesting to analyse the relationship of RL and R1 in additional arabidopsis ecotypes and genotypes to test further how leaf number affects branching in arabidopsis. Future research should also address the role of the FT-mediated and other flowering pathways on branching and how sugar and Tre6P signalling might interact with the flowering pathway during this process.

The correlation analyses of the different ecotypes grown in different photoperiods confirmed the strong correlation (close to R^2^ = 1) of C1 and CL (Fig. 6, 7, S6, S7). This highlights that there is almost no plasticity in cauline branching per se, with every cauline leaf giving rise to 1 cauline branch (Fig. 8A). This is in stark contrast to rosette branching which correlated only weakly with rosette leaf number when long-day grown ecotypes were analysed separately (Fig. 2D, S1) showing that branching at rosette nodes is not simply a consequence of leaf number and is potentially regulated by the integration of many other endogenous and exogenous signals (Fig. 8A).

In contrast to rosette buds, cauline buds might receive different signals because of their location on the main stem. Due to this position, they are continuously exposed to red light potentially resulting in very low levels of BRC1 and ABA (González-Grandío et al., 2013; Reddy et al., 2013; Yao and Finlayson, 2015; González-Grandío et al., 2017; Holalu and Finlayson, 2017). This might be the cause of the strong activation of cauline branches and might be a potential reason why cauline buds behave differently from rosette buds in terms of activation and outgrowth. Future studies should aim at addressing these differences in cauline and rosette bud outgrowth in detail and would also benefit from determining the extent to which axillary buds may form different numbers of leaves prior to rapid elongation into a mature branch (Ferguson and Beveridge, 2009; Barbier et al., 2019).

## CONCLUSIONS ON PHENOTYPING SHOOT BRANCHING

We show that C1 and R1 are rarely negatively correlated in arabidopsis. Therefore, in our view, accurate phenotyping of shoot branching in arabidopsis should show C1 and R1 separately, and interpretations should not be based on the total number of primary branches (Fig. 8B). Cauline branching is highly correlated to the number of cauline nodes produced, which in turn is related, to a large extent, to flowering time. We highlight here that the mechanism controlling rosette branching involves not only hormonal and nutrient (including sugar) signalling pathways, but also involves flowering regulation, light signalling and potentially further unknown signalling pathways. In highly branched strigolactone mutants, RL and R1 are highly correlated variables. In this case, RL as well as R1/RL are useful to distinguish small genetic and environmental effects on shoot branching as well as independent effects on branching in plants that flower differently (Fig. 8B).

## MATERIAL AND METHODS

### Plant material and growth

Branching and flowering data from different laboratories working on branching in arabidopsis were collected and used in this study. *Arabidopsis thaliana* Columbia-0 (Col-0), Landsberg *erecta* (Ler) or Wassilewskija (Ws-4) ecotypes and mutants in these backgrounds were used. Some parts of the data were published previously, including the Fig. 1 experiment (Exp) 3 (Aguilar-Martínez et al., 2007), Fig. 1 Exp 5 (Brewer et al., 2016), Fig. 4A/5A Exp 1 and Exp 2 (Fichtner et al., 2021b). Arabidopsis plants were all grown in a 16-h photoperiod unless otherwise stated according to the following light and temperature conditions: All plants from condition A were grown on UQ23 potting mix (70% composted pine bark 0–5mm, 30% coco-peat) supplemented with dolomite and osmocote, using light intensities of 120 to 150 μmol m^-2^ s^-1^ (unless otherwise stated) and a temperature of 22°C day/ 18°C night (except for experiment in higher temperature in Fig. 4B). All plants from condition B were grown in a 1:1 mixture of soil (Stender) and vermiculite using light intensities of 150 μmol m^-2^ s^-1^ and a temperature of 22°C day/ 18°C night. All plants from condition C were grown in Seed & Cutting Premium Germinating mix (Debco) at 23°C constant temperature and 75 μmol m^-2^ s^-1^ (C.1) or 120 μmol m^-2^ s^-1^ (C.2) light intensity. All plants from condition D were grown as described in Aguilar-Martínez et al. (2007) using light intensities of 120 μmol m^-2^ s^-1^ and a temperature of 20°C. All plants from condition E were grown at different densities (1, 3 or 10 plants per 33 cm^2^ cell) on a mixture of three parts seed and modular compost plus sand (Scott Levington) to one part vermiculite for horticultural use (Sinclair), at a light intensity of 240 μmol m^-2^ s^-1^ and a temperature of 23°C. In the case of Fig. S4A the different arabidopsis ecotypes were all grown in the same cabinets using the same soil type (condition A) but under a large variety of different photoperiods and light regimes. Different photoperiods and light regimes included: black, 16-h photoperiod with 150 μmol m^-2^ s^-1^ light intensity; grey, 16-h photoperiod with 4 weeks of 150 μmol m^-2^ s^-1^ and a subsequent shift to 40 μmol m^-2^ s^-1^ light intensity; blue, 16-h photoperiod with 75 μmol m^-2^ s^-1^ light intensity; green, 4 weeks in an 8-h photoperiod then shift to a 16-h photoperiod (150 μmol m^-2^ s^-1^ light intensity each); yellow, 8-h photoperiod with 150 μmol m^-2^ s^-1^ light intensity; red, 8 hours of 150 μmol m^-2^ s^-1^ light intensity followed by 8 hours of 5 μmol m^-2^ s^-1^ light intensity; light blue (Col-0 only), 18-h photoperiod with 150 μmol m^-2^ s^-1^ light intensity.

### Arabidopsis mutant lines

All arabidopsis mutant lines used in this study were described earlier: *brc1* mutants (Aguilar-Martínez et al., 2007); *lbo-10* (*lbo-1* mutation backcrossed six times to Col-0, thus termed here *lbo-10*) and *lbo-1, max4-9* and *lbo-1 max4-9* (Ws-4 background) (Rasmussen et al., 2012; Brewer et al., 2016); *d27-1* (Waters et al., 2012); *max1-1* and *max2-1* (Stirnberg et al., 2002); *max3-9* (Booker et al., 2004); *max4-1* (Sorefan et al., 2003); *d14-1* (Chevalier et al., 2014); *htl-3* (a *kai2* allele isolated in Col-0) (Toh et al., 2014); *smxl678* (*smxl6-4,7-3,8-1*) with the *max2-1* mutation crossed out (Soundappan et al., 2015); *35S:YUCCA1* (also referred to as *yuc1D*) (Zhao et al., 2001); *ft-10* and *35S:FT* (Yoo et al., 2005); *soc1-6* (Wang et al., 2009); *fd-3* (Searle et al., 2009).

### Phenotyping

Rosette and cauline leaves were counted separately to give rosette leaf (RL) and cauline leaf (CL) numbers. Primary rosette (R1) and cauline (C1) branches (shoots ≥ 0.5 cm) were counted and R1 and C1 were added to give the total primary branch number (T1). RL and R1 were used to determine primary rosette branch number per leaf (R1/RL).

### Statistical analysis and data visualization

Data analyses and plotting were performed using R Studio Version 1.4.1717 (www.rstudio.com) with R version 4.1.0 (https://cran.r-project.org/) and the packages ggplot2, stats and agricolae using Pearson’s correlation or an ANOVA based post hoc comparison of means test (Fisher’s least significant difference (LSD) test). Heatmap analyses were performed with the heatmap.2 function (R package heatmaply) using the agglomeration method “average” for the hierarchical cluster analysis of genotypes, correlation-based clustering of traits and the distance measure “canberra” for the computation of the distance matrix. Principal component analyses were done using the R package factoextra. Figures were compiled using Adobe Illustrator 2021.

## ACKNOWLEDGEMENTS

We thank Cecilia Corben and Ursula Krause for excellent technical assistance and help with plant growth and phenotyping. We thank Mark Stitt for helpful comments on the manuscript. Fig. 1A and Fig. 8 were created with BioRender.com. This work was supported by the Australian Research Council (F.F.B., C.A.B., Discovery grant DP150102086; F.F., F.F.B, C.A.B., Georgina Sweet Laureate Fellowship FL180100139 to C.A.B., and Future Fellowship FT180100081 to P.B.B.), the Max Planck Society (F.F.), and BIO2014-57011-R, BIO2017-84363-R and FEDER funds (P.C.).

