## Supplementary Information for "Different plasticity of bud outgrowth at cauline and rosette nodes in *Arabidopsis thaliana*"

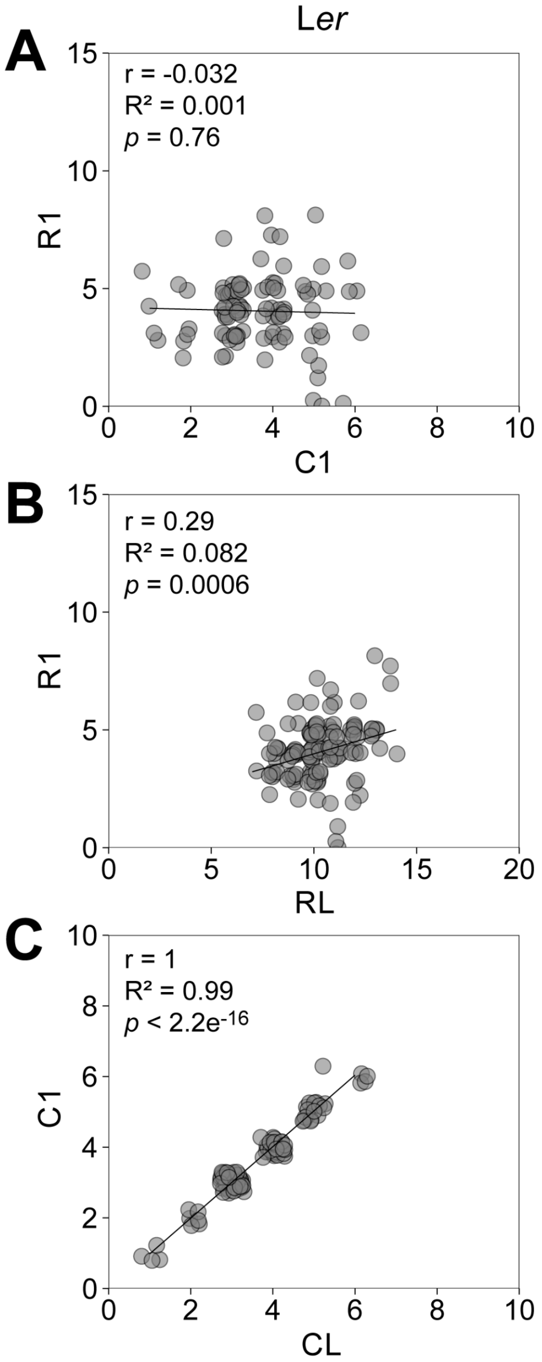


**Figure S1.** Correlation analysis of Landsberg *erecta* (L*er*) wild-type plants. (A) The number of primary cauline branches (C1) was correlated against the number of rosette branches (R1). (B) The number of rosette leaves (RL) was correlated against R1. (D) The number of C1 was correlated against the number of and cauline leaves (CL). The Pearson correlation coefficient (r), coefficient of determination (R2) and probability (p) were calculated. All plants were grown in a 16-h photoperiod. Genotypes are indicated by different colours. Each data point represents a single plant. Data points were jittered to avoid overplotting and were alpha blended meaning that regions of high point density show up as areas of high colour intensity.

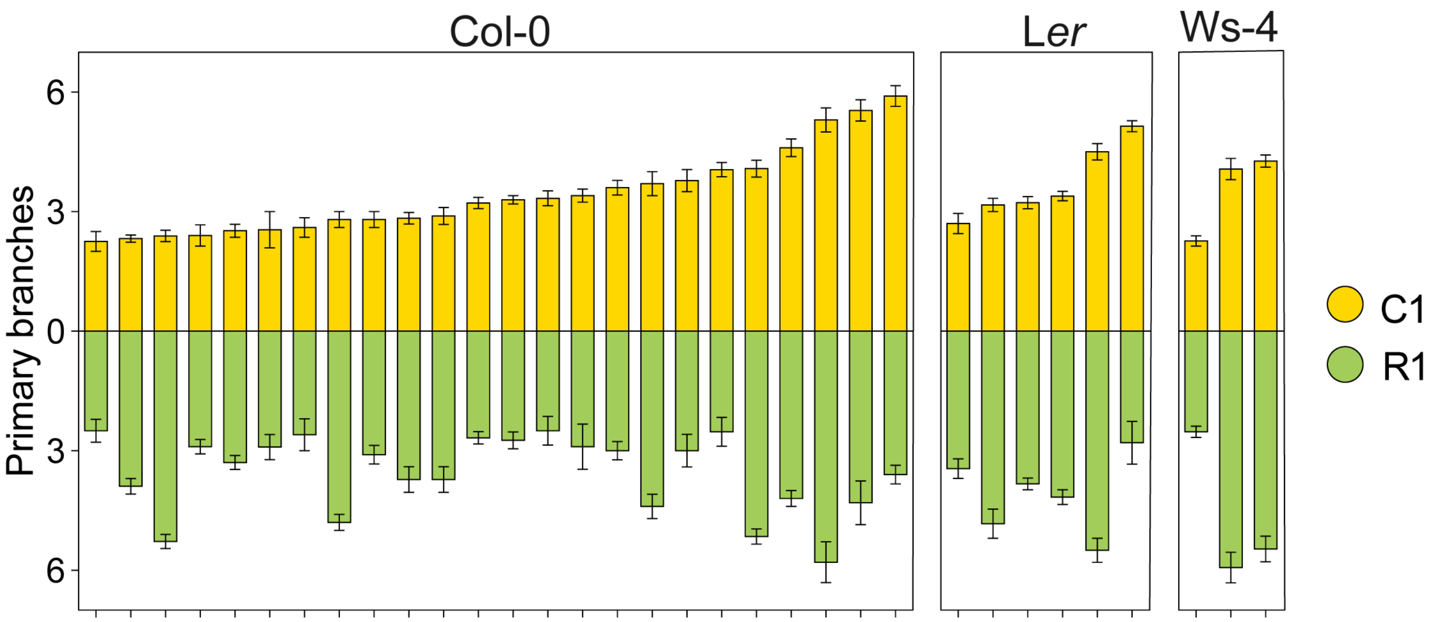


**Figure S2.** The variation in cauline and rosette branching is different in three arabidopsis ecotypes. Arabidopsis wild types (Columbia-0, Col-0; Landsberg *erecta*, L*er*; Wassilewskija-4, Ws-4) were grown in 16-h photoperiods and normal planting densities. Cauline (yellow, C1) and rosette (green, R1) branch numbers are plotted separately and in ascending order for the number of C1. Depicted is the mean ± SEM.

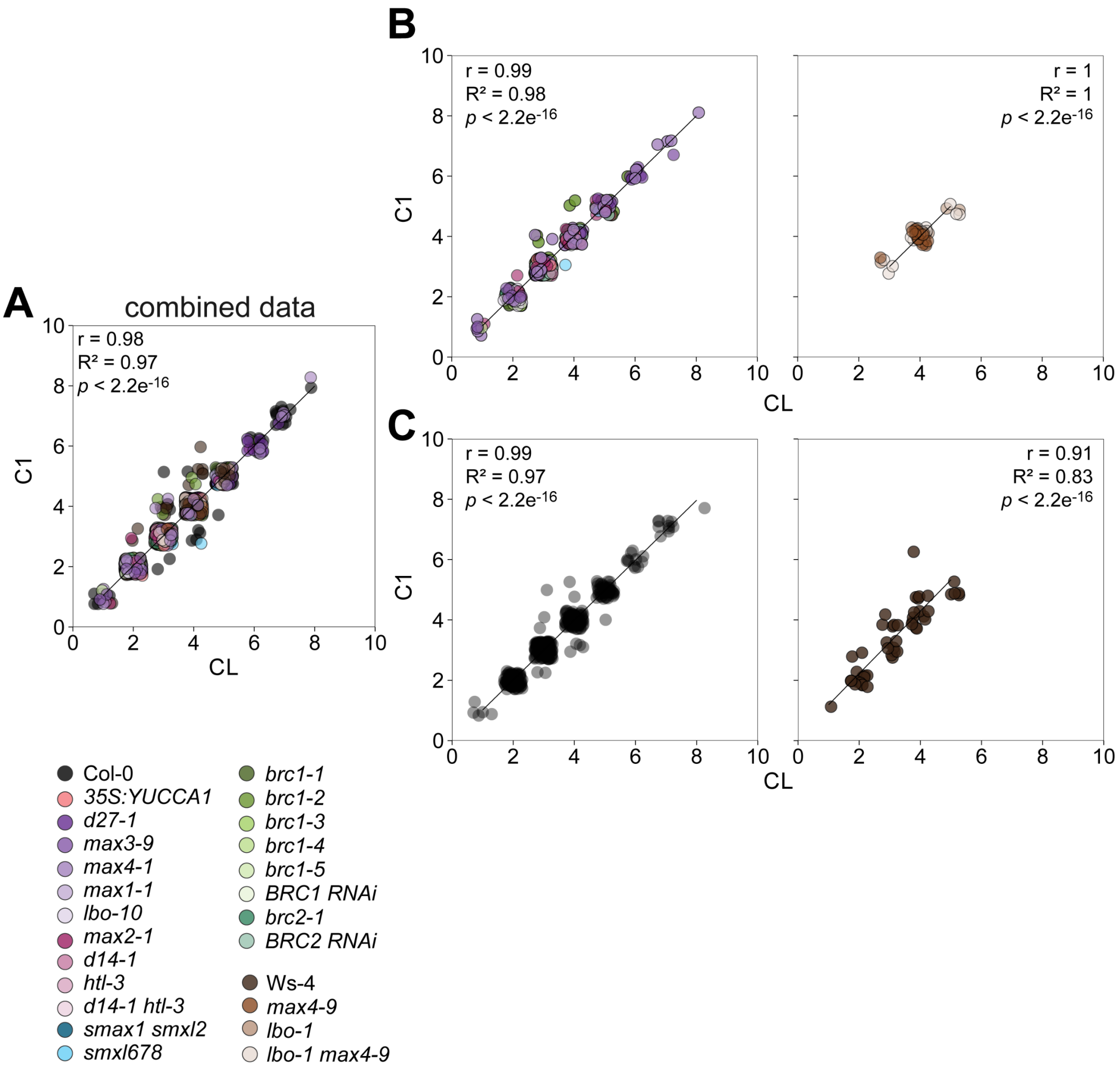


**Figure S3.** Correlation analysis of the number of cauline leaves (CL) and cauline branches (C1) in arabidopsis wild type and mutant plants. The Pearson correlation coefficient (r), coefficient of determination ($R^2$) and probability (*p*) were calculated. (A) The correlation for all data presented in Figure 2 was used. (B) The correlations of CL and C1 were separated by the corresponding ecotype. (C) The correlation of CL and C1 for wild-type plants only. All plants were grown in a 16-h photoperiod. Genotypes are indicated by different colours. Each data point represents a single plant. Data points were jittered to avoid overplotting and were alpha blended meaning that regions of high point density show up as areas of high colour intensity.

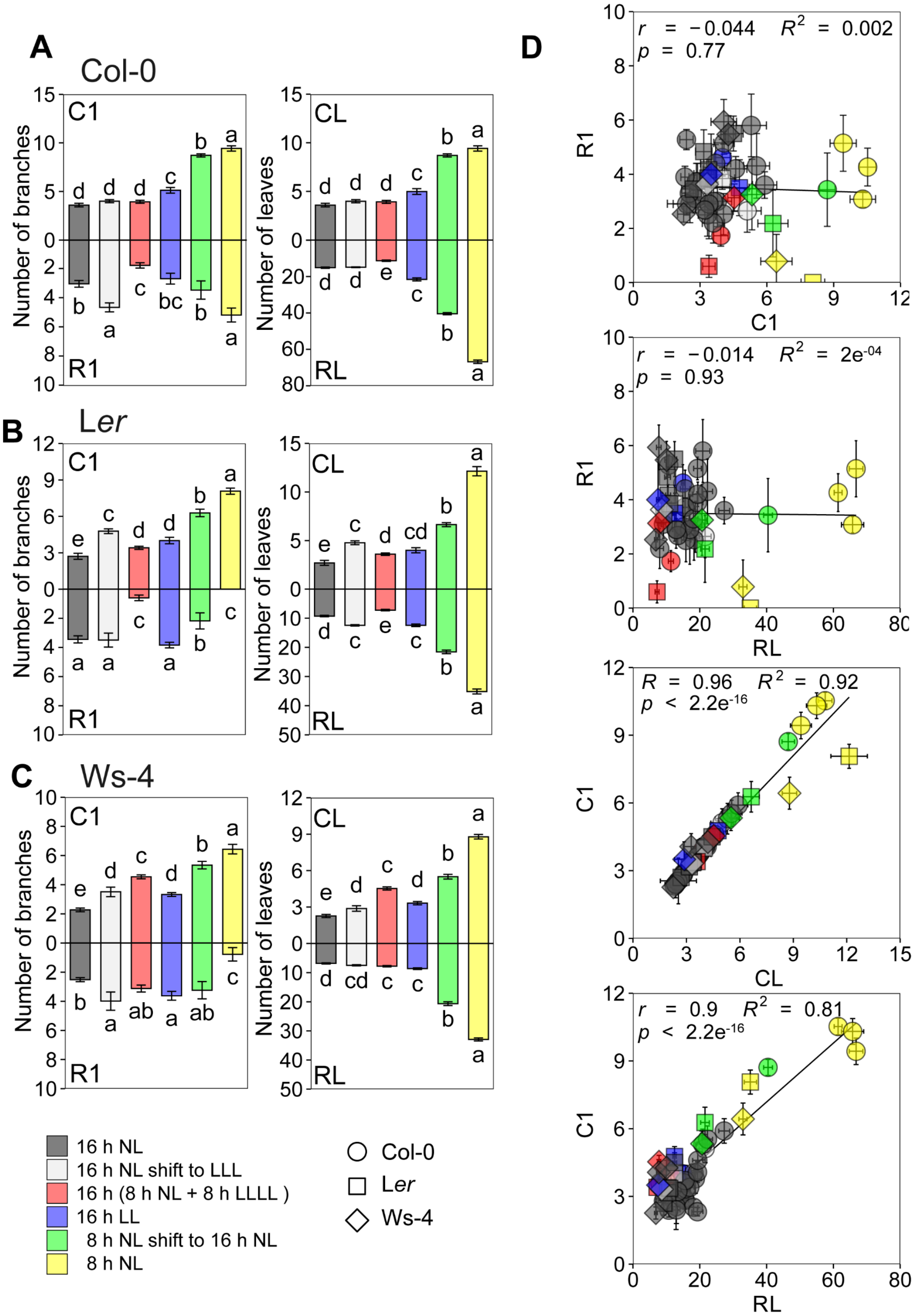


**Figure S4.** Different arabidopsis ecotypes (Columbia-0, Col-0 (circles); Landsberg *erecta*, Ler (squares); Wassilewskija-4, Ws-4 (diamonds)) were grown in a variety of photoperiods as indicated in Figure 6A. (A) The number of cauline and rosette branches (C1 and R1, respectively) and leaf numbers (CL and RL, respectively) were determined. A single experiment for each photoperiod and ecotype that was done under the same conditions is shown as an example. Letters represent significant differences based on ANOVA with post-hoc LSD testing ($p < 0.05$). Depicted is the mean ± SEM. (B) Correlation analyses of C1 and R1, R1 and rosette leaves (RL), C1 and cauline leaves (CL), and C1 and RL. The Pearson correlation coefficient (r), coefficient of determination ($R^2$) and probability ($p$) were calculated based on the mean values for each experiment. Each data point represents an independent experiment. Data points were alpha blended meaning that regions of high point density show up as areas of high colour intensity.

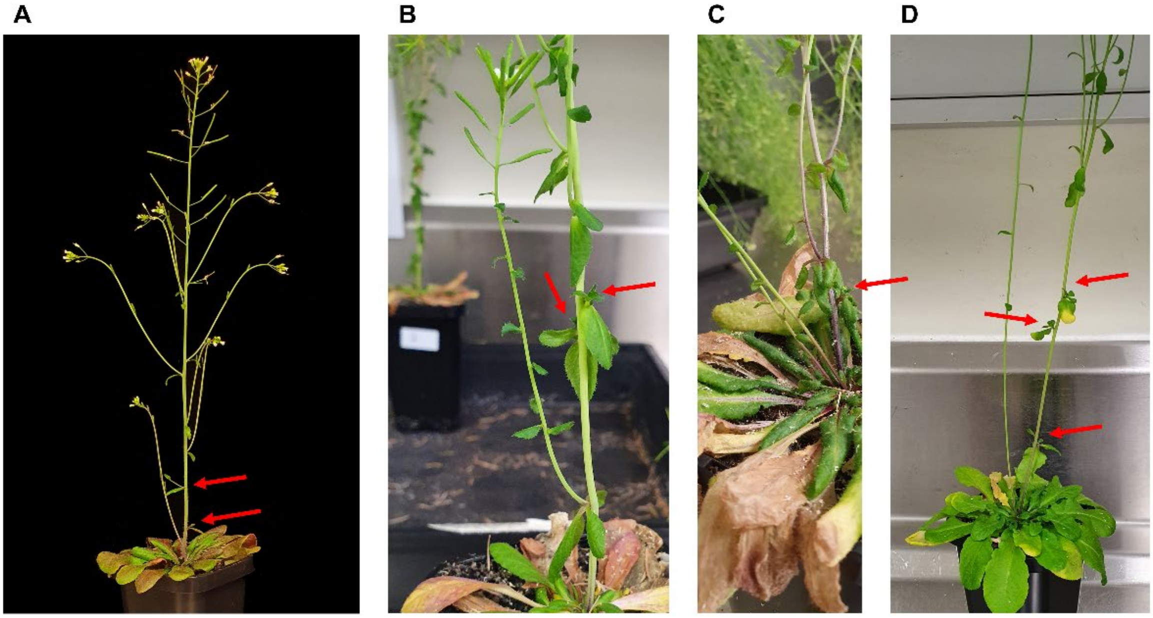


**Figure S5.** Branch skipping phenomenon in different arabidopsis ecotypes observed in short day or short day to long day shift conditions. Primary cauline branches that did not elongate are marked by red arrows. Lower node cauline branches are elongating before upper node cauline branches in arabidopsis (A) Columbia-0 (Col-0) plants that were grown for 4 weeks in an 8-h photoperiod before shifted to a 16-h photoperiod, and in (B) Landsberg *erecta* plants grown in an 8-h photoperiod. Rosette branches are elongating before upper node cauline branches in arabidopsis (C) Col-0 plants and (D) Wassilewskija-4 plants grown in an 8-h photoperiod. This phenomena was not observed when plants were grown in long days.

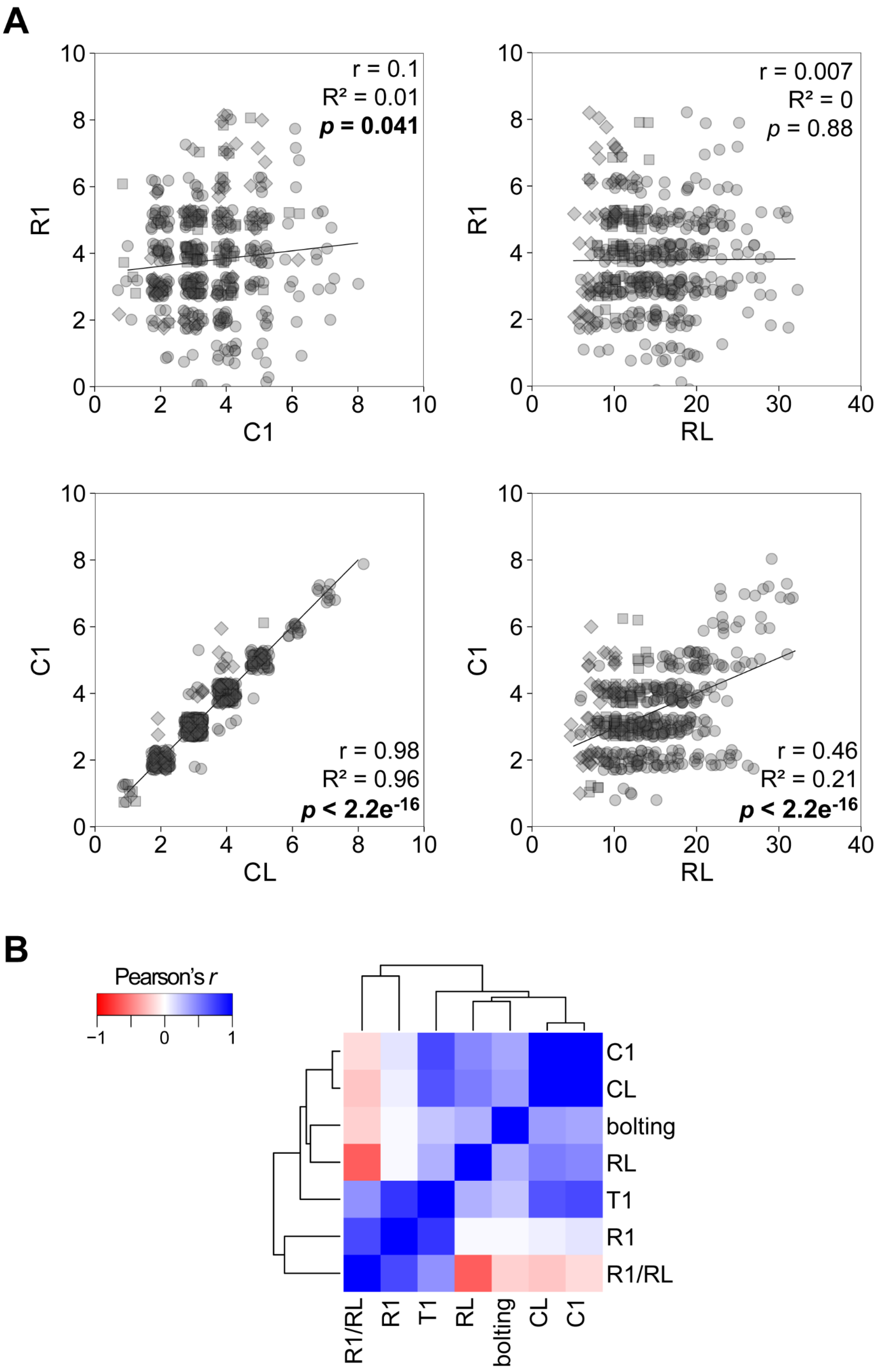


**Figure S6.** Different arabidopsis ecotypes were grown in long-day photoperiods. Correlation analyses of primary cauline branches (C1) and rosette branches (R1), R1 and rosette leaves (RL), C1 and cauline leaves (CL), and C1 and RL in different arabidopsis ecotypes (Columbia-0 (circles); Landsberg *erecta* (squares); Wassilewskija-4, Ws-4 (diamonds)). The Pearson correlation coefficient (r), coefficient of determination ($R^2$) and probability (*p*) were calculated based on the mean values. Each data point represents an independent experiment. Significant correlations are indicated in bold. Data points were jittered to avoid overplotting and were alpha blended meaning that regions of high point density show up as areas of high colour intensity. (B) Heatmap analysis of the Pearson's r of branching and flowering traits. Dendrograms represent clusters based on a Canberra distance matrix with average-based clustering. bolting, days to bolting.
